# The structure of *Mycobacterium tuberculosis* heme-degrading protein, MhuD, in complex with product

**DOI:** 10.1101/731950

**Authors:** Alex Chao, Kalistyn H. Burley, Paul J. Sieminski, David L. Mobley, Celia W. Goulding

**Affiliations:** Department of Molecular Biology & Biochemistry, University of California Irvine, Irvine, CA 92697; Pharmaceutical Sciences, University of California Irvine, Irvine, CA 92697; Chemistry, University of California Irvine, Irvine, CA 92697

**Keywords:** Heme degradation, *Mycobacterium tuberculosis*, mycobilin, biliverdin, iron

## Abstract

*Mycobacterium tuberculosis* (Mtb), the causative agent of tuberculosis, requires iron for survival. In Mtb, MhuD is the cytosolic protein that degrades imported heme. MhuD is distinct, both in sequence and structure, from canonical heme oxygenases (HOs) but homologous with IsdG-type proteins. Canonical HO is found mainly in eukaryotes, while IsdG-type proteins are predominantly found in prokaryotes including pathogens. While there are several published structures of MhuD and other IsdG-type proteins in complex with heme substrate, no structures have been reported of IsdG-type proteins in complex with product, unlike HOs. We recently showed that the Mtb variant MhuD-R26S produces biliverdin IXα (αBV) rather than the wild-type (WT) mycobilin isomers as product. Given that mycobilin and other IsdG-type protein products like staphylobilin are difficult to isolate in quantities sufficient for structure determination, here we use the MhuD-R26S variant and its product αBV as a proxy to study the IsdG-type protein/product complex. First we show that αBV has nanomolar affinity for MhuD and the R26S variant. Second we determined the MhuD-R26S-αBV complex structure to 2.5 Å, which reveals two notable features (1) two αBV molecules bound per active site and (2) a new α-helix (α3) as compared with the MhuD-heme structure. Finally, by molecular dynamics simulations we show that α3 is stable with the proximal αBV alone. MhuD’s high affinity for its product and structural and electrostatic changes that accompany substrate turnover suggest that there is an unidentified protein that is responsible for product extraction from MhuD and other IsdG-type proteins.

## Introduction

Heme degradation is important for a variety of biological functions, including iron re-utilization, cell signaling, and antioxidant defense (Brouard et al., 2000; Dore et al., 1999; Ferris et al., 1999). The well-studied canonical heme oxygenase (HO), human HO (hHO- 1), catalyzes the oxidative cleavage of heme to release biliverdin XIα (αBV, Figure 1A), ferrous iron, and carbon monoxide (CO) (Matsui et al., 2010; Tenhunen et al., 1969; Yoshida et al., 1980). HO homologs have also been found in many eukaryotes and some prokaryotes including the pathogens, *Corynebacterium diphtheriae*, *Pseudomonas aeruginosa* and *Neisseria meningitidis*, and predominately produce the same heme degradation products (Ratliff et al., 2001; Schmitt, 1997; Wilks, 2002; Wilks and Schmitt, 1998; Zhu et al., 2000). In eukaryotes, the HO reaction is coupled with the conversion of biliverdin (BV) to bilirubin by biliverdin reductase (BVR) (Noguchi et al., 1979). After conjugation of bilirubin with glucuronic acid, bilirubin is excreted (Mantle, 2002). The fate of HO-produced BV in prokaryotes has not been well studied thus far. Although in *P. aeruginosa*, it has been shown that extracellular acquired heme is degraded by a HO homolog, HemO, whereby the BV by-product is then excreted, but not further reduced, by an unknown mechanism (Barker et al., 2012).

**Figure 1.**
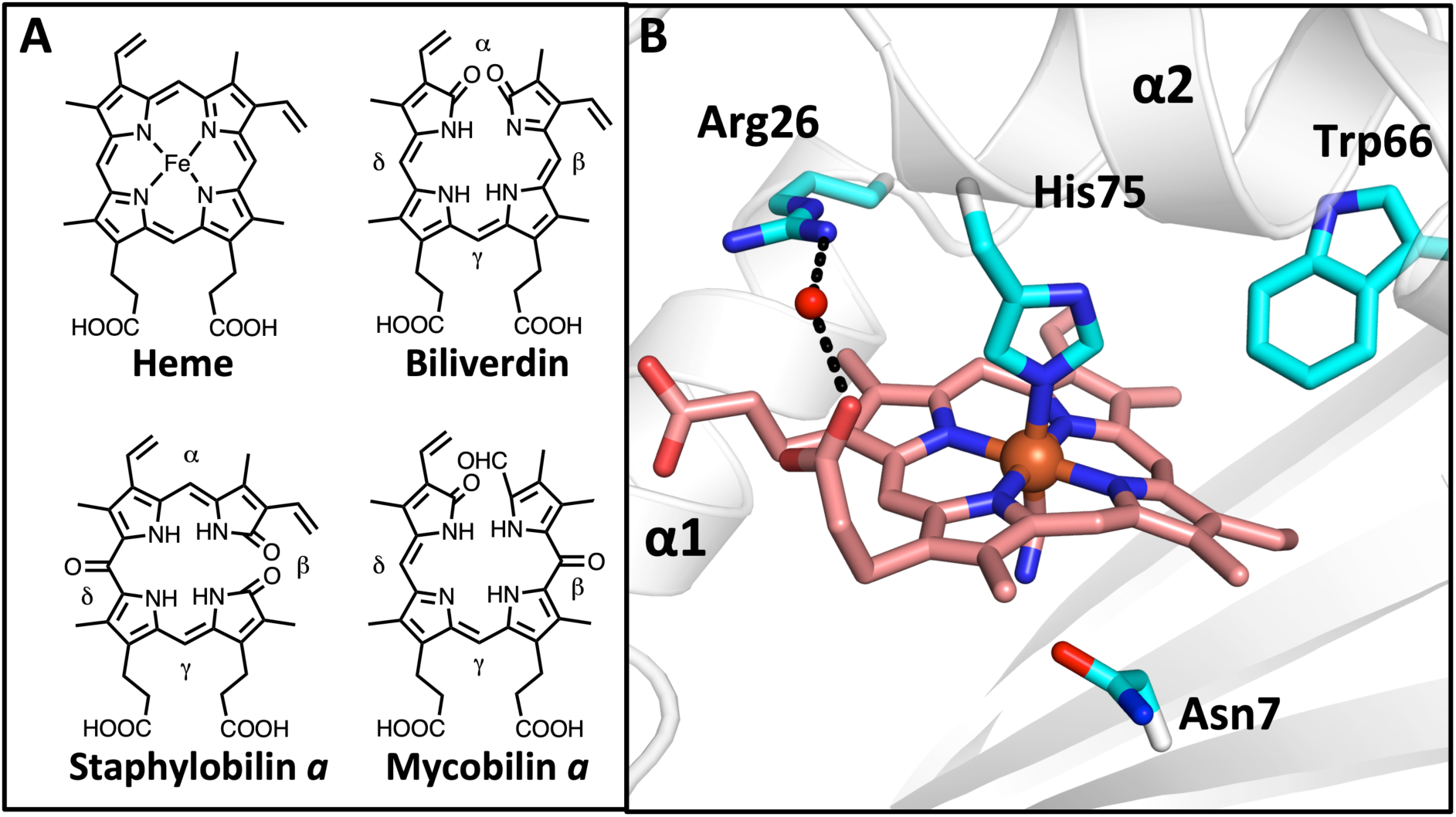
Structures of tetrapyrroles and WT-MhuD-monoheme. **A.** The structure of heme and heme tetrapyrrole degradation products. **B**. The structure of the active site of the Mtb heme-degrading protein, MhuD, in its cyano-derivatized monoheme form (cartoon, white). Depicted in stick representation (cyan) are essential Asn7, Trp66 and His75 (which coordinates heme-iron) residues, and Arg26 that forms a water-mediated H-bond with one of the heme propionates.

There is a heme-degrading protein family distinct, both in sequence and structure, from canonical HOs mainly found in bacteria, and termed the *i*ron *s*urface *d*eterminan*t* G (IsdG)-type family (Chim et al., 2010; Skaar et al., 2004; Wu et al., 2005). *Staphylococcus aureus* IsdG and IsdI were the first members characterized (Skaar et al., 2004; Wu et al., 2005), and instead of degrading heme to BV, iron and CO, these enzymes cleave and oxidize heme at the β- and δ-meso carbon sites to produce staphylobilin isomers (Figure 1A), free iron and formaldehyde (Matsui et al., 2013; Reniere et al., 2010). Other IsdG- type enzymes have been identified including MhuD from *Mycobacterium tuberculosis* (Mtb) and LFO1 from eukaryotic *Chlamydomonas reinhardtii*, and both degrade heme into unique products (Chim et al., 2010; Lojek et al., 2017). While LFO1 degrades heme into a yet-to-be-determined product(s) (Lojek et al., 2017), MhuD has been shown to degrade heme into iron and mycobilin isomers (Figure 1A) (Nambu et al., 2013). Like staphylobilins, the mycobilin isomers are also oxidized at the β- or δ-meso carbons; however cleavage occurs at the α-meso carbon with no observed loss of a C1-product (Nambu et al., 2013). It was proposed that there is no C1-product as hHO-1 produced CO triggers the transition of Mtb from its active to latent state (Nambu et al., 2013). The fate of the IsdG-type protein heme degradation tetrapyrrole products, like staphylobilin and mycobilin, is unknown; however they may have antioxidant properties similar to BV (Vanella et al., 2016).

The structures of HO and IsdG-type proteins are quite distinct. HOs are comprised of a monomeric α-helical domain (Schuller et al., 1999), while IsdG-type proteins consist of a dimeric *β*-barrel decorated with two α-helices from each monomer (Wu et al., 2005). Unsurprisingly, the two distinct classes of heme-degrading enzymes degrade heme by different mechanisms (Matsui et al., 2016). Although heme is coordinated by a proximal His residue in both HO and IsdG-type enzymes, the heme molecule in HOs is near-planar, while for IsdG-type proteins, the heme is ruffled (Schuller et al., 1999; Wu et al., 2005). Furthermore, HO has a distal pocket with a network of ordered water molecules that facilitates the three consecutive monooxygenase steps required for heme degradation (Matsui et al., 2010). In contrast, the distal heme pocket is quite hydrophobic for IsdG- type proteins, with only one or two ordered waters observed in the proximal pocket (Graves et al., 2014; Wu et al., 2005). In MhuD, it has been proposed that the hydrophobic pocket together with the ruffled heme contributes to the sequential monooxygenase and dioxygenase steps necessary for MhuD to degrade heme (Matsui et al., 2016).

The structures of both human and *C. diphtheriae* HO complexed with αBV illustrate that the HO heme degradation reaction is coupled with a conformational change from a ‘closed’ to ‘open’ state (Lad et al., 2004; Unno et al., 2013). In both the eukaryotic and prokaryotic HO structures, the open-product bound state results from relaxation of the distal and proximal helices, and a rotation of the catalytic His side-chain out of the active site pocket as it is no longer coordinated to heme-iron (Lad et al., 2004; Unno et al., 2013). This structural shift suggests that a degree of protein flexibility is necessary amongst the HO homologs to facilitate catalysis. Structures of apo and monoheme bound forms of IsdG-type proteins reveal an analogous conformational change in the catalytic His residue as it is absent or disordered in the apo structures; however the ordering of the elongated L2 loop region upon heme binding is a much more drastic conformational shift as compared with HOs (apo-MhuD; Protein Data Bank (PDB) ID: 5UQ4) (Graves et al., 2014; Lee et al., 2008; Wu et al., 2005). Compared with other studied IsdG-type proteins, the MhuD active site is exceptionally flexible and can accommodate two molecules of heme, resulting in protein inactivation (Chim et al., 2010; Graves et al., 2014). The biological significance of this conformational plasticity and its role in product turnover is not well understood as there is no structure of an IsdG-type enzyme in its product-bound form.

The structure of an IsdG-type protein in complex with its heme degradation product would further our understanding of the mechanism of action of this protein family. Staphylobilin and mycobilin products of IsdG-type proteins are difficult to purify (Nambu et al., 2013; Reniere et al., 2010), which has presumably hindered the structural analysis of the product-bound form. Recently, we demonstrated that a MhuD variant, MhuD-Arg26Ser (Figure 1B), upon heme degradation produces αBV, formaldehyde and iron (Chao and Goulding, 2019). In this current study, we determine the affinity of MhuD and the MhuD- R26S variant to both heme and αBV, and show they both bind heme and αBV in the nanomolar range. This high affinity to αBV has allowed us to utilize the MhuD-R26S variant as a proxy to study the IsdG-type proteins in complex with product. Upon solving the crystal structure of the MhuD-R26S-αBV, we observed the formation of a novel secondary structural element as compared to the heme-bound MhuD structure, and its implications will be discussed further.

## Methods

### Fluorescence-detection of ligand binding

Fluorescence-detected titrations of heme and αBV were carried out using a protocol previously described (Thakuri et al., 2018). Stock solutions of MhuD (80 nM), heme (8 µM), and αBV (8 µM) were prepared in 50 mM Tris/HCl pH 7.4, 150 mM NaCl. Heme or αBV was titrated and gently mixed in 16 nM or 32 nM increments into MhuD-WT or MhuD- R26S. Following a 1-min incubation, fluorescence emission spectra were acquired between 320 to 500 nm on a Hitachi F-4500 Fluorescence Spectrophotometer through excitation at 285 nm with the following parameters: 1/3 nm step size, scan speed of 240 nm/min, PMT voltage of 700 V, and slit widths at 10 nm (MhuD WT) and 20 nm (MhuD- R26S).

### Fluorescence emission spectral analysis

Results from the above fluorescence-based assay were fitted to Eqn. 1 derived from Conger et el (Conger et al., 2017; Thakuri et al., 2018), to determine the equilibrium dissociation-constant (*K_d_*) of heme or αBV with MhuD and its mutants.

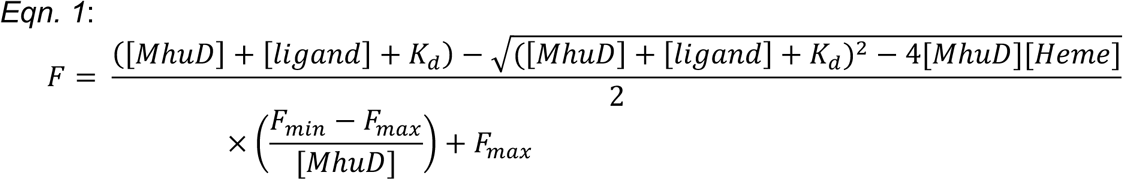

In Eqn. 1, [MhuD] is the total concentration of MhuD or mutant MhuD, [ligand] is the total concentration of heme or αBV, F_max_ is the emission intensity with ligand absent, and F_min_ is the emission intensity for fully ligand-bound MhuD. Fitting of the fluorescence emission intensity at 340 nm for *K_d_* determination was performed using Origin 2018.

### Expression and purification of Mtb MhuD and the R26S variant

Wild-type (WT) MhuD and the R26S variant were purified as previously reported (Chao and Goulding, 2019; Chim et al., 2010). In brief, *E. coli* B21-Gold (DE3) cells transformed with pET22b-MhuD plasmid were grown in LB medium containing 50 μg/mL ampicillin at 37°C. Overexpression was induced at OD_600_ of ∼ 0.6 using 1 mM IPTG. The cells were harvested 4 hours post induction and resuspended in lysis buffer (50 mM Tris/HCl pH 7.4, 350 mM NaCl and 10 mM imidazole). Cells were lysed via sonication and the resulting lysate was centrifuged at 14,000 rpm. The cell supernatant was loaded onto a Ni^2+^- charged HiTrap chelating column (5 mL) and washed with lysis buffer. Bound protein was eluted from the column with increasing concentrations of imidazole. Next, MhuD, which elutes at 50 and 100 mM imidazole, was concentrated (Amicon, 5 kDa molecular mass cutoff) and was further purified on a S75 gel filtration column in 20 mM Tris/HCl pH 8, and 10 mM NaCl.

### Crystallization of MhuD-R26S-αBV complex

To prepare an αBV solution, approximately 2 mg of αBV hydrochloride (SigmaAldrich) was dissolved in 500 μL 0.1 M NaOH followed by 500 μL 1 M Tris/HCl pH 7.4 before dilution into 50 mM Tris/HCl pH 7.4, 150 mM NaCl. A ferric chloride solution was prepared by dissolving 27.3 mg of ferric chloride hexahydrate (SigmaAldrich) in 10 mL water. A 1.3 fold excess of a 1:1 molar ratio solution of αBV and ferric chloride was gradually added to 100 μM apo-MhuD R26S and incubated overnight at 4°C before being concentrated to 10 mg/mL (Lowry assay (Lowry et al., 1951)). Light blue crystals appeared in 0.1 M HEPES pH 6.5, 4.6 M NaCl, 30 mM glycyl-glycyl-glycine after 2 days. The crystals were flash frozen in 100% NVH oil and a data set to 2.5 Å was collected at 70K. The collected data was indexed, integrated, and reduced using iMOSFLM (Battye et al., 2011). Initial phase determination was carried out using Phaser in the PHENIX suite (Adams et al., 2010) using WT MhuD-heme-CN structure as a search model (PDB ID 4NL5) (Graves et al., 2014). αBV molecules were positioned into appropriate positive electron density in the vicinity of the active site, and the structure was refined using phenix.refine (Adams et al., 2010). For each promoter, the electron density of the loop region between Ala24 to Asn32 is poorly defined and residues His25-Val30 were modeled as alanines. The only Ramanchandran outliers are Val30 modeled as Ala in both Chain A and B, which are in this poorly defined region of electron density.

### Molecular dynamics (MD) simulations

The input files for MhuD^4NL5^-heme and MhuD^4NL5^-αBV simulations were prepared from the structure of dimeric MhuD-heme-CN and MhuD^4NL5^-heme did not include the cyano group (PDB ID: 4NL5). For the MhuD-αBV complex, input files were prepared from the MhuD-R26S-αBV structure, wherein the R26S mutation was reversed and just one αBV (proximal) was retained per active site. All crystallographic waters were removed with the exception of ordered waters in the active site of MhuD^4NL5^ (HOH numbers: 313, 325, 349). Hydrogen atoms were added to the system using pdb4amber from AmberTools15 with default protonation states (Case et al., 2015). For simulations of MhuD^4NL5^-αBV, the coordinates of heme atoms served as a scaffold upon which each αBV was manually docked. αBV was parameterized using antechamber from AmberTools15 (Case et al., 2015) with GAFF version 1.7 and AM1-BCC charges. The ligated His75-Fe-heme ligand was parameterized according to previously published Density Functional Theory calculations (Harris et al., 2001). Missing atoms for each structure were added using tleap in AmberTools15 (Case et al., 2015), and parameterized using ff99SBildn (Lindorff-Larsen et al., 2010). Each system was explicitly solvated in tleap with a 10 Å rectangular box of TIP3P water, extending from the surface of the protein to the box edge, and sodium ions were added to neutralize the charge of the system.

Minimization proceeded using *sander* from Amber14 (Case et al., 2015) with steepest descents running for 20,000 timesteps of 2 fs each, followed by heating from 100 K to 300 K with constant volume for 25,000 timesteps of 2 fs. Equilibration continued using *sander* for 500,000 timesteps of 2 fs under constant pressure with positional restraints initially applied on all non-water atoms at 50 kcal/mol/Å^2^ and progressively lifted in increments of 5 kcal/mol/Å^2^ over ten 50,000 step segments. The resulting topology and coordinate files for each system were used as inputs for MD simulations. Production simulations were executed in OpenMM 7.1.0 (Eastman et al., 2017) using a Langevin integrator with a 2 fs timestep and a friction coefficient of 10/ps. For each of the three systems, five independent simulations of 100 ns each were initiated with randomized velocities. Among the five simulations, one simulation was restarted for each system for an additional 500 ns, bringing the total simulation time to 1 μs/system.

### MD analysis

Distances and secondary structure assignments were computed using MDTraj (McGibbon et al., 2015). For analysis, each monomer of MhuD was treated independently with the assumption that long-range interactions between each subunit are negligible for the time scales simulated here. To analyze the orientation of His75 during simulations, two orientations were defined: 1) *Active Site* – side chain pointing into binding pocket, towards the center of the tetrapyrrole, and 2) *Solvent Exposed* – side chain flipped away from the binding pocket, as in the MhuD-αBV structure (see Figure 4Bi). To assign these positions, we computed the distance between the *ε* nitrogen atom (furthest from the backbone) on His75 and the nitrogen atom located between the α and β carbons on either αBV or heme. If the distance was less than 6 Å, the His75 residue was classified as being oriented in the *Active Site* position; otherwise the position was classified as *Solvent Exposed*.

For the Arg79 positional analysis, we identified four positions; 1) *Active site* – Arg79 directed into the binding pocket interacting with the ligand, 2) *Helix 1* – Arg79 side-chain interacting with residues on α-helix-1 (i.e residues 16-25) of MhuD, 3) *Helix 2* – Arg79 side-chain interacting with residues on α-helix-2 (residues 60-75), and 4) *Solvent Exposed* – Arg79 side-chain oriented into the surrounding solvent. To analyze the Arg79 in our simulations, position assignments were defined as follows: 1) *Active Site* – guanidinium carbon of Arg79 side-chain (CZ) within 4.5 Å of either terminal carbon on the propionate groups of the heme or αBV ligands. 2) *Helix 1 -* Arg79 CZ atom within 6 Å of Glu16 α carbon 3) *Helix 2* – Arg79 CZ atom within 6 Å of His75 or Ile72 backbone oxygen and more than 4.5 Å from terminal carbons on the propionate groups of the heme or αBV ligands. 4) *Solvent Exposed* – Any position of Arg79 falling outside the description of positions 1-3.

## Results

### Affinity of substrate and product to MhuD

Previously, the affinity (*K*_d_) of WT MhuD for heme was measured to be ∼6 nM (Thakuri et al., 2018); empirically it has been also been observed that MhuD also has a high affinity for its product, as isolation of MhuD mycobilin products requires an iron-chelator followed by protein denaturation (Chao and Goulding, 2019; Nambu et al., 2013). Recently, we demonstrated that the MhuD-R26S variant degrades heme to produce αBV as its predominate tetrapyrrole product rather than the WT MhuD mycobilin isomers (Chao and Goulding, 2019; Nambu et al., 2013). As it is difficult to isolate large quantities of mycobilin, we use the MhuD-R26S variant and its αBV product as a model system to study IsdG-type proteins in complex with product. First, we determined the affinities of heme and αBV to both WT MhuD and the R26S variant. The heme affinity was measured using a previously described fluorescence-based assay (Figure 2) (Thakuri et al., 2018). The *K*d of heme for MhuD-WT and MhuD-R26S are similar, both about 6 nM (Table 1), whereas the *K*d values of αBV for MhuD-WT and MhuD-R26S are ∼36 nM and ∼92 nM, respectively (Table 1). The fact that WT MhuD binds αBV with a higher affinity than the R26S variant (even though the product of WT MhuD is mycobilin rather than αBV) implies that the Arg26 residue may form contacts with the tetrapyrrole product.

**Figure 2.**
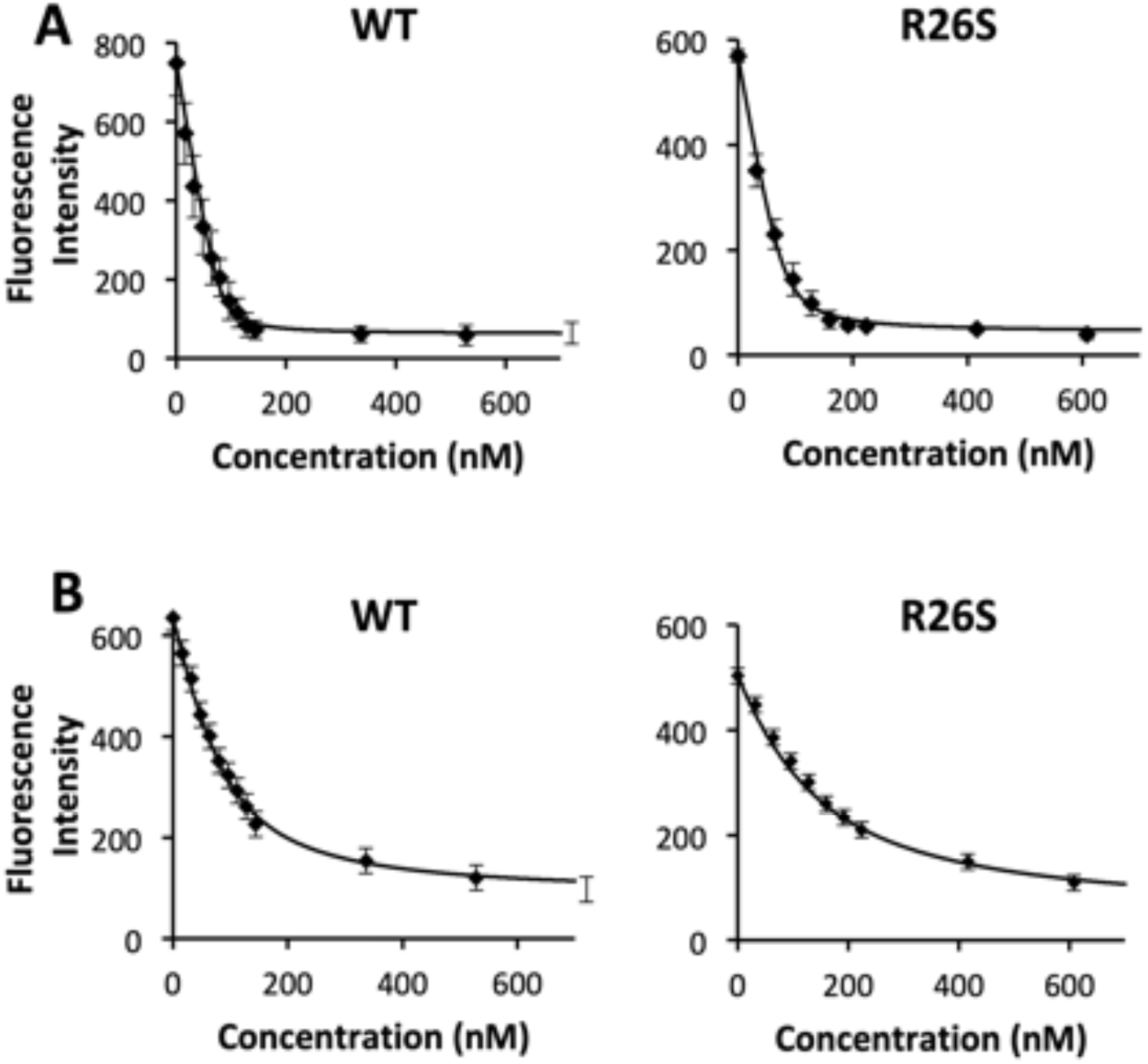
Heme and αBV affinity of MhuD and the R26S variant. Representative fluorescent emission intensities at 340 nm after excitation at 280 nm of WT MhuD (left panels) and the MhuD-R26S variant (right panels) with increasing concentrations of **A.** heme and **B.** αBV.

**Table 1.** Heme and αBV binding affinities (*K_d_*) were determined for WT MhuD and the R26S variant complexes with 1:1 heme or αBV to protein ratio. Each experiment was performed in triplicate.

| | heme $K_d$ (nM) | αBV $K_d$ (nM) |
| --- | --- | --- |
| MhuD-WT | 6.0 ± 2.9 | 36.4 ± 2.1 |
| MhuD-R26S | 9.5 ± 3.0 | 92.4 ± 6.8 |

### The structure of the MhuD-R26S-αBV complex

To investigate the structural impacts on MhuD arising from heme degradation into product, we turned our attention to structure determination of the MhuD-R26S variant in complex with αBV. The MhuD-R26S-αBV complex crystallized in the presence of ferric chloride, and light blue crystals appeared in 0.1 HEPES pH 6.5, 4.6 M NaCl and 30 mM glycyl-glycyl-glycine after 2 days. The structure of the MhuD-R26S-αBV complex was solved to 2.5 Å resolution with a final R/R_free_ of 23.2/28.5 (Table 2). The asymmetric unit contains two MhuD-R26S promoters bridged between their active sites by five partially π- stacked αBV molecules (Figure 3A). The MhuD-R26S monomers each complexed with two αBV molecules superimpose with a root-mean-square deviation (RMSD) of 0.2 Å. The biologically relevant MhuD-R26S homodimer is observed in the crystallographic 2- fold symmetry (Figure 3B), similar to the apo- and heme-bound MhuD structures (apo-MhuD; PDB code: 5UQ4) (Chim et al., 2010; Graves et al., 2014).

**Figure 3.**
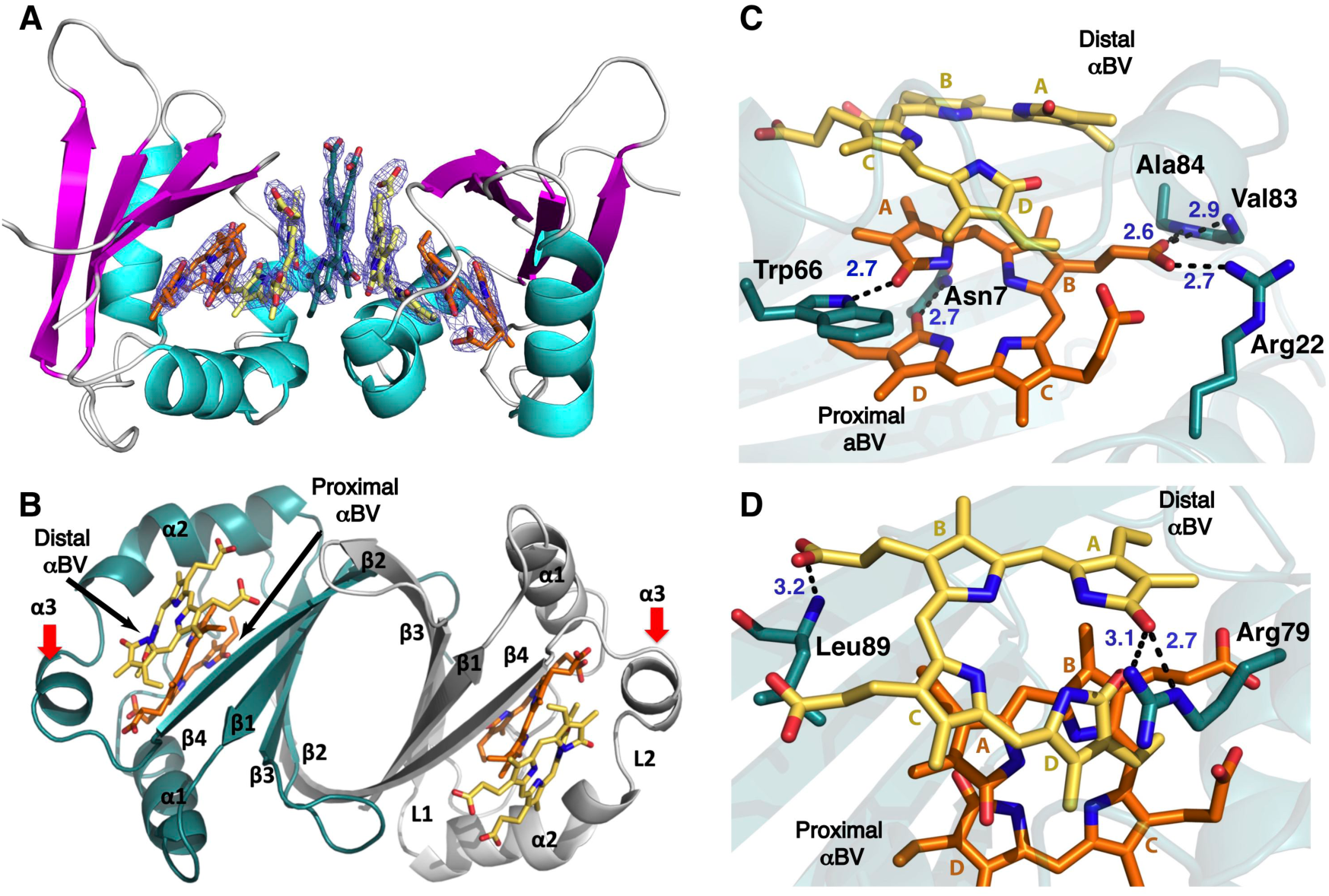
Structure of the MhuD-R26S-αBV complex. **A.** The asymmetric unit that contains five molecules of αBV stacked connecting the active sites of two MhuD monomers. **B.** The dimeric biological assembly with two molecules of αBV per active site. The new structural helix, α3, is denoted with a red arrow. **C.** Interactions of the proximal αBV (orange) with MhuD (green). Blue dashed lines represent H-bonds with their length in Å. **D.** Interactions of the distal αBV (yellow) with MhuD (green).

**Table 2.** X-ray diffraction data and atomic refinement for MhuD-R26S-αBV complex.

|  |  |
| --- | --- |
| Space Group | I222 |
| Unit cell dimensions (Å) | 37.22 x 113.61 x 113.65 |
| pH of crystallization condition | 6.5 |
| Protein concentration (mg/mL) | 20 |
| **Data Collection** |  |
| Wavelength, Å | 1.0 |
| Resolution range | 35.94 – 2.5 |
| Unique reflections (total) | 8713 (17426) |
| Completeness, %* | 99.89 (100.00) |
| Redundancy* | 2.0 (2.0) |
| $R_{\text{merge}}^{\ast, \dagger}$ | 0.013 (0.062) |
| $CC_{1/2}^{\ast}$ | 1.0 (0.99) |
| $I/\sigma^{\ast}$ | 25.41 (10.64) |
| NCS copies | 2 |
| **Model refinement** |  |
| Resolution range, Å | 35.93 - 2.5 |
| No. of reflections (working/free) | 8707 (867) |
| No. of protein + ligand atoms | 1430 |
| No. of water molecules | 16 |
| No. of αBVs in NCS | 5 |
| $R_{\text{work}}/R_{\text{free}}$, %<sup>¶</sup> | 23.16/28.53 |
| **Rms deviations** |  |
| Bond lengths, Å | 0.009 |
| Bond angles | 1.52 |
| **Ramachandran plot** |  |
| Most favorable region, % | 90.96 |
| Additional allowed region, % | 7.98 |
| Outliers (Val30), % | 1.06 |
| **PDB ID code** | **6PLE** |

\*Statistics for the highest-resolution shell are given in parentheses.
<sup>†</sup> $R_{\text{merge}} = \sum_{hkl} \sum_i |I_i(hkl) - \langle I(hkl) \rangle| / \sum_{hkl} \sum_i I_i(hkl)$
<sup>¶</sup> $R_{\text{work}} = \sum |F_{\text{obs}} - F_{\text{calc}}| / \sum F_{\text{obs}}$ . $R_{\text{free}}$ was computed identically except where all reflections belong to a test set of 10% randomly selected data.

The MhuD-R26S-αBV structure is similar to that of the MhuD-heme-CN and MhuD- diheme structures (Chim et al., 2010; Graves et al., 2014), where each subunit forms a ferredoxin-like fold. Two monomers form a dimeric antiparallel β-barrel where each monomer has three α-helices together with two flexible loop regions; the first loop connects α-helix-1 to β-strand-2 and the second connects α-helix-3 to β-strand-4 (Figure 3B). It should be noted that the topology of the MhuD-R26S-αBV complex differs from the structures of apo-MhuD and heme-bound WT-MhuD (Graves et al., 2014). Within the MhuD-R26S-αBV structure, there is an additional short α-helix-3 formed by residues Ala76-Asn81, which was not observed in the monoheme, diheme and apo-MhuD structures (Chim et al., 2010; Graves et al., 2014), and thus the latter structures have a slightly longer extended L2 loop region. There are also other subtle conformational changes that will be discussed later.

### αBV Binding to MhuD

Each MhuD-R26S active site binds two molecules of αBV (Figures 3B). The solvent-protected or proximal αBV interacts with both MhuD and the distal αBV, and the distal αBV also interacts with MhuD. The proximal αBV forms five electrostatic and eleven hydrophobic interactions with MhuD residues. The αBV propionate-6 carboxylate group hydrogen-bonds (H-bonds) with the backbone amides of Val83 and Ala84 (2.9 and 2.6 Å, respectively), and forms a salt-bridge with the NH1 group of Arg22 (2.7 Å). Additionally, Trp66 NE1 H-bonds (2.7 Å) with the A-ring lactam oxygen (=O), and Asn7 ND2 H-bonds (2.7 Å) to the D-ring lactam oxygen (Figure 3C). The tetrapyrrole plane of the distal αBV is nearly parallel to that of the proximal αBV.

The proximal and distal αBV molecules interact through hydrophobic and van der Waals interactions only. The distal αBV is rotated ∼205° and flipped ∼180° with respect to the αBV tetrapyrrole plane, and the distal αBV tetrapyrrole plane is positioned ∼3.4 Å above and translated ∼5.0 Å relative to the proximal αBV plane. Consequently, the C-ring of the distal αBV is partially placed over the A-ring of the proximal αBV resulting in the distal αBV being more solvent exposed than the proximal one (Figure 3D). The proximal αBV forms five H-bonds with MhuD whereas the distal αBV forms only three and has fewer hydrophobic interactions with MhuD than the proximal αBV (Figure 3D). The A-ring lactam oxygen of the distal αBV H-bonds with Arg79 NE and NH1 (3.1 and 2.7 Å, respectively), and the distal αBV propionate-6 H-bonds to the Leu89 amide group (3.2 Å).

Along with the proximal and distal αBV molecules, there is a third ‘bridging’ αBV, which connects the two symmetrical promoters each bound to two αBV molecules (Figure 3A). The third ‘bridging’ αBV is parallel to the tetrapyrrole plane of the flanking distal αBV molecules from each monomer and is separated by ∼3.8 Å from each distal αBV. Additionally, the bridging αBV is rotated ∼90° and translated ∼2.5 Å with respect to the distal αBV molecules. The bridging αBV interacts with the distal αBVs by hydrophobic and van der Waals interactions, and also forms one H-bond to each, where the bridging αBV ring-D/A lactam oxygen H-bonds to the ring-A/D pyrrole nitrogen of the distal αBVs (2.8 Å and 3.3 Å), respectively. Finally, the bridging αBV forms a H-bond to each MhuD promoter, ring-A/D lactam oxygen H-bonds with Arg79 NH2 group on respective monomers (3.0 and 2.6 Å, respectively).

### Comparison of MhuD substrate and product bound structures

There are several distinct structural differences between the inactive substrate-(MhuD- heme-CN) (Graves et al., 2014) and product-bound (MhuD-R26S-αBV) MhuD structures, despite an RMSD of 1.1 Å (Figure 4A). The most notable difference between the two structures is within α-helix-2 and sequential loop region L2. In MhuD–heme–CN, the α- helix-2 is kinked after residue Asn68 and terminates at His75, where this region in the MhuD-R26S-αBV structure has a looser helical geometry. This kinked helical region in the MhuD–heme–CN structure positions the catalytic His75 within the active site so it coordinates with heme-iron. However, in MhuD-R26S-αBV, His75 flips out of the active site (90° rotation and translation of 2.8 Å of the Cα, Figure 4Bi) and is stabilized by a H-bond between the His75 imidazole nitrogen to the backbone carbonyl of Ala27 (3.2 Å). In combination with the slight unraveling of the C-terminus of α-helix-2 in MhuD-R26S- αBV, the extended loop L2 region has an additional α-helix (α3, Ala76-Asn81) as compared to MhuD–heme–CN (Graves et al., 2014) (Figure 4A). Within the L2 loop region of MhuD–heme–CN, both His78 and Arg79 are solvent-exposed with no observable electron density for the Arg79 side-chain. In contrast, in MhuD-R26S-αBV, Arg79 is located in α-helix-3, and its side-chain is positioned towards the active site and is stabilized by a H-bond between its NH1 group with the backbone carbonyl of Ile72 (3.1 Å) (Figure 4Bii); notably His78 is still solvent-exposed and weakly stabilized by a cation-π interaction between its imidazole side-chain and Arg22 (3.7 Å).

**Figure 4.**
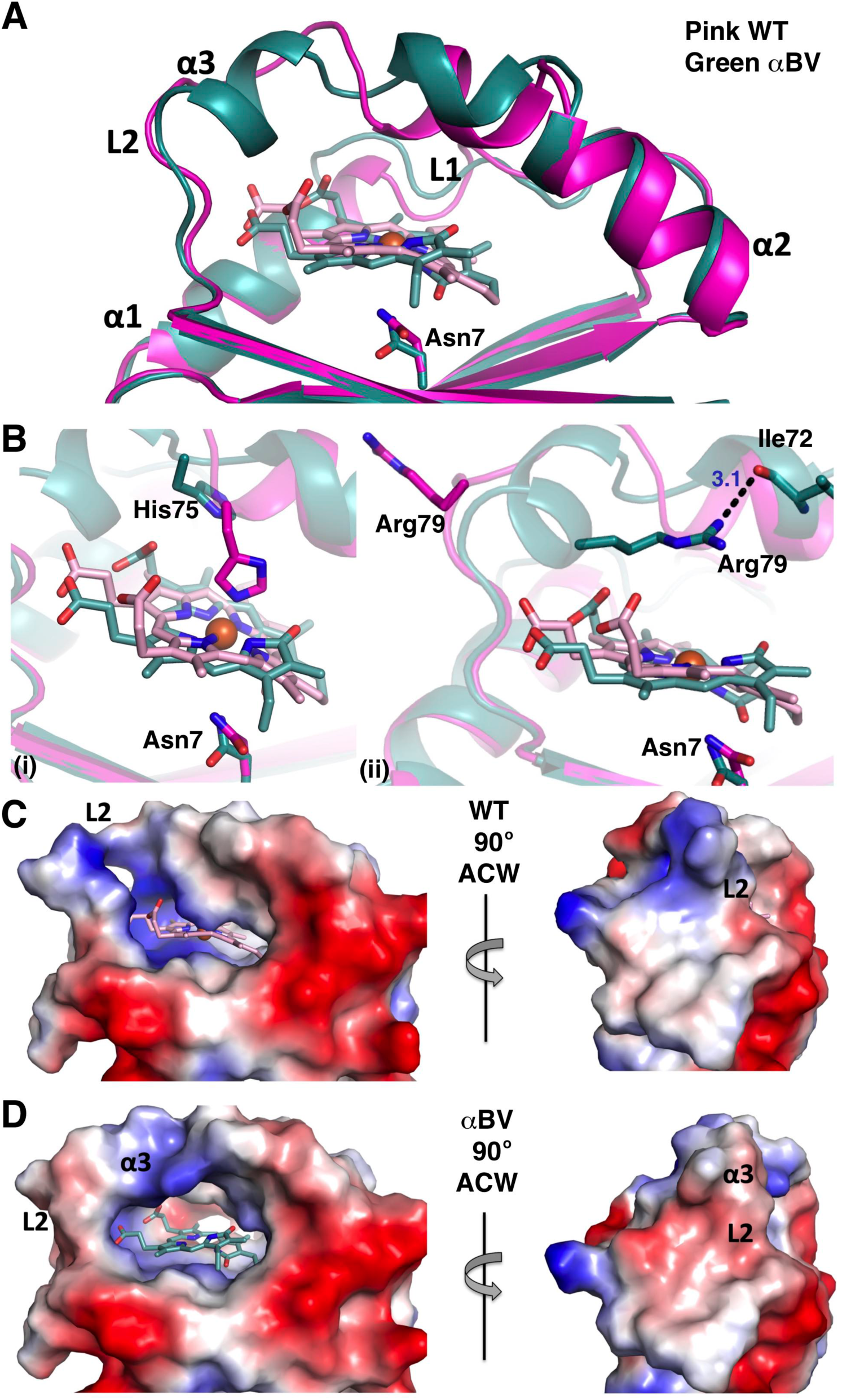
Structural comparison of MhuD-heme-CN and MhuD-R26S-αBV. **A.** Superposition of the MhuD-heme-CN (pink, PDB ID 4NL5, the cyano group is omitted for clarity) and MhuD-R26S-αBV (green). **B**. Active site comparison with **(i)** catalytic residues Asn7 and His75, and **(ii)** residues Asn7 and Arg79, are shown in stick representation. Black dashed lines represent H-bonds with their length in Å. **C-D** Electrostatic molecular surface representations, where blue and red are positively and negatively charged, respectively. Right panel is the left panel rotated 90**°** anticlockwise (ACW). **C.** MhuD- heme-CN (WT) and **D.** MhuD-R26S-αBV (αBV).

Another minor structural difference between substrate and product bound MhuD complexes is the unraveling of the C-terminal α-helix-1 in MhuD-R26S-αBV compared to MhuD-heme-CN. Arg26 is situated in this location and participates in a water-mediated H-bond with one of the heme propionates in the MhuD-heme-CN structure (Graves et al., 2014) (Figures 1B & 4A). Within the MhuD-R26S-αBV structure, this region is no longer helical. Thus, this observed unraveling of α-helix-1 may be the result of the Arg26Ser mutation, however, this conformational change ensures that the backbone carbonyl group of the subsequent residue, Ala27, is in H-bonding distance of the imidazole nitrogen of His75, to stabilize the flipped out His75 in the MhuD-R26S-αBV structure (Nambu et al., 2013).

The conformational change from going from substrate to product bound also alters the active site pocket volume and the electrostatic potential of the molecular surface (Figures 4C & 4D). The active site volume increases dramatically from heme bound to that of αBV, from 188 to 354 Å^3^, calculated utilizing CASTp (Tian et al., 2018). In conjunction with the increased active site volume, the molecular surface region surrounding the active site and the L2 loop region undergoes an electrostatic potential change. The MhuD molecular surface surrounding one side of the exposed active site is positively and negatively charged in the presence of heme (Figure 4C), whereas in the presence of αBV it becomes more hydrophobic and negatively charged with a positively charged bridge capping the active site (Figure 4D). Furthermore, when rotated 90° about the y-axis, there is an altered molecular surface electrostatics from positively charged and hydrophobic to predominately negatively charged in the presence of heme and αBV, respectively (Figures 4C & 4D, right panels).

### Proximal αBV is representative of the MhuD physiological product

Within the MhuD-R26S-αBV structure, the orientation of the proximal αBV is representative of the MhuD physiological product, mycobilin. The proximal αBV adopts a similar orientation as heme in the MhuD-heme-CN structure (Figures 4A-B) (Graves et al., 2014), however αBV is rotated approximately 20° about the plane compared to heme-CN and the tetrapyrrole ring structure of αBV is considerably more twisted compared to heme. Additionally, the overall conformation of MhuD proximal αBV is similar to the αBV product of *C. diphtheriae* HmuO (PDB 4GPC) (Unno et al., 2013), representative of the all-Z-all-syn type BV conformation. However, in another structure of hHO-1 in complex with αBV (PDB 1S8C) (Lad et al., 2004), the αBV occupies an internal cavity adjacent to the active site and exhibits a more linear extended conformation. This was proposed to be the route of αBV dissociation from hHO-1 in the absence of BVR (Lad et al., 2004). As αBV is bound tightly to MhuD, we did not anticipate that we would observe a partially dissociated product conformation or location, as seen in the hHO-1 structure. Thus these observations suggest that the proximal αBV within MhuD-R26S-αBV structure is the physiological orientation of the MhuD tetrapyrrole product, mycobilin, within the active site.

### MD Simulations

MD simulations were performed to gain further insight into the protein conformational changes associated with heme and αBV binding. In particular, we wanted to evaluate the *in silico* stability of the MhuD α-helix-3 (Ala76-Asn81) in the presence of proximal αBV alone. MD simulations were set up using the biological dimer of the MhuD-R26S-αBV structure with the R26S mutation modeled as WT Arg26 and only the proximal αBVs retained. Over the 1 μs of combined simulation time, including a continuous 600 ns simulation, the α-helix-3 persisted for over 70% of the run, suggesting that the α-helix-3 is stable when there is just one αBV molecule present per active site. Interestingly, we also observe some helix formation within the L1 loop region, an otherwise highly flexible region of the protein (Figure 5A).

**Figure 5.**
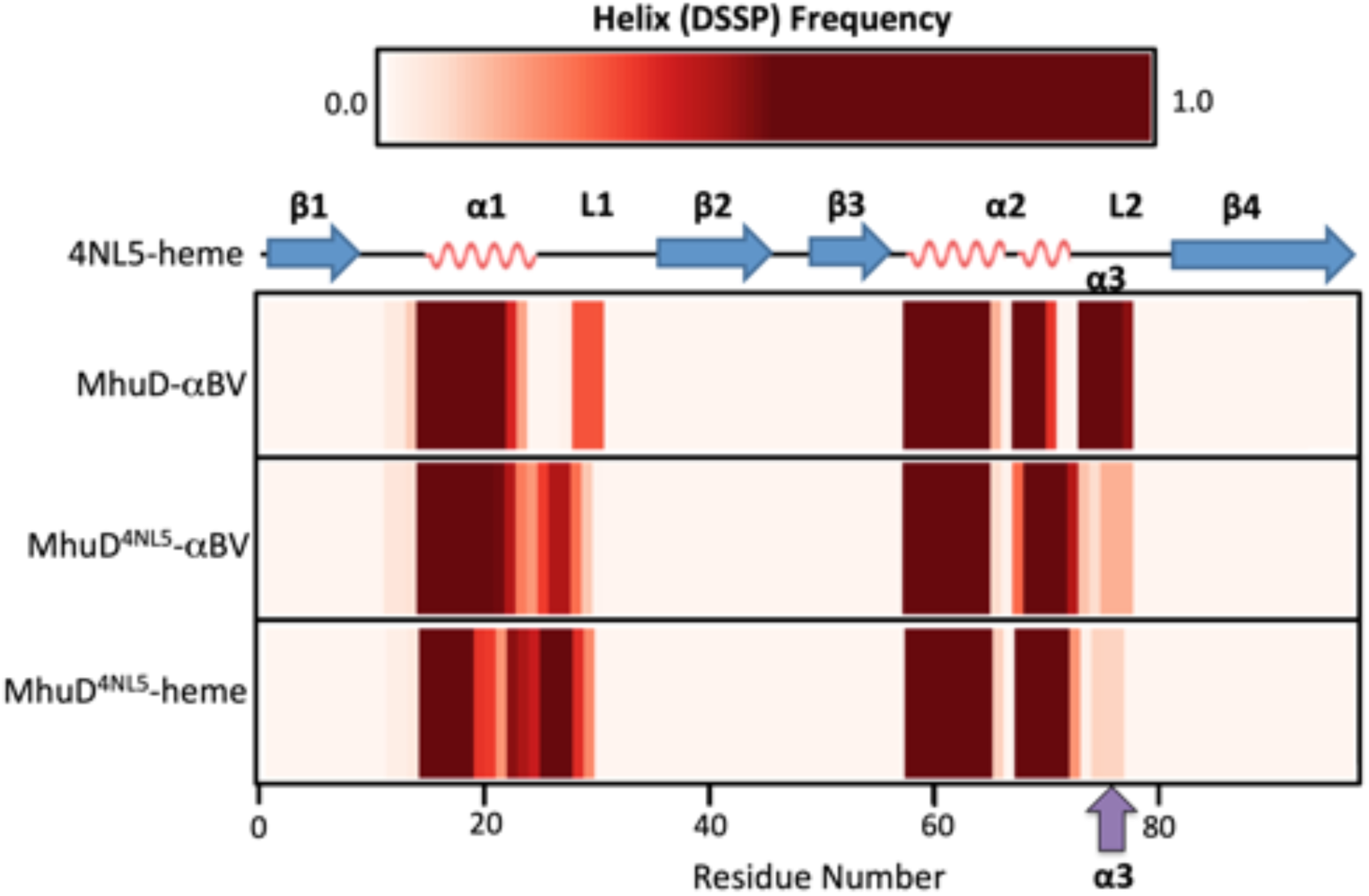
Helical stability in MD simulations of MhuD. The frequency of helix formation (DSSP) is plotted for each residue of MhuD in simulations with αBV and heme. For simulations of MhuD-αBV initiated from the coordinates of the MhuD-αBV structure, the novel α-helix (α3) that forms in L2 persists. In simulations of MhuD^4NL5^-αBV initiated from the coordinates of the MhuD-monoheme structure (PDB: 4NL5), α3 transiently forms in the L2 region. In the presence of heme (MhuD^4NL5^-heme), formation of α3 also is observed but less frequently than in MhuD^4NL5^-αBV.

To test if the α-helix-3 forms during turnover from heme to product, two more sets of simulations were carried out. The first contained the MhuD-heme-CN structure without the cyano group (MhuD^4NL5^-heme) while the second was comprised of the MhuD^4NL5^ protein structure with αBV docked in place of heme to give MhuD^4NL5^-αBV. As with MhuD- αBV, these simulations each ran for a total of 1 μs. Although the two αBV MD systems (MhuD-αBV and MhuD^4NL5^-αBV) are identical in composition, they have distinct initial positions and velocities; MhuD-αBV simulations start from the coordinates of the αBV- bound crystal structure while MhuD^4NL5^-αBV simulations start from those of the heme-bound structure. Given sufficient simulation time, the dynamics of these two systems should eventually converge. While we did not reach convergence, it is compelling that we see some *de novo* formation of the α-helix-3 in our simulation of MhuD^4NL5^-αBV (Figure 5B), suggesting that it is a relevant structural motif that forms in the presence of αBV. We suspect that the α-helix-3 may be further stabilized by contact of MhuD with other interacting proteins. This α-helix-3 is also transiently observed in the MhuD^4NL5^-heme simulation (Figure 5C); however its occurrence is less frequent compared to MhuD^4NL5^- αBV.

We also sought to evaluate whether the 90° rotation of the catalytic His75 side chain away from the active site, as observed in the MhuD-αBV structure (Figures 4Bi & 6), is facilitated by substrate turnover to αBV. In the MhuD-αBV simulations, His75 residue remains in the “flipped out” or solvent exposed orientation (Figure 6B) while in the MhuD^4NL5^-heme simulations, it is ligated to the heme-iron atom and thus remains tethered to the active site. For MhuD^4NL5^-αBV, the His75 alternates between the two orientations (Figure 6C), as shown in Figures 6A & 6B. The solvent exposed position may be further stabilized in the MhuD^4NL5^-αBV simulations after extended simulation time and upon full formation of the α-helix-3, as observed in the MhuD-αBV structure, where the Arg79 side-chain forms a hydrogen bond with the backbone carbonyl of Ile72 (Figure 4Bii).

**Figure 6.**
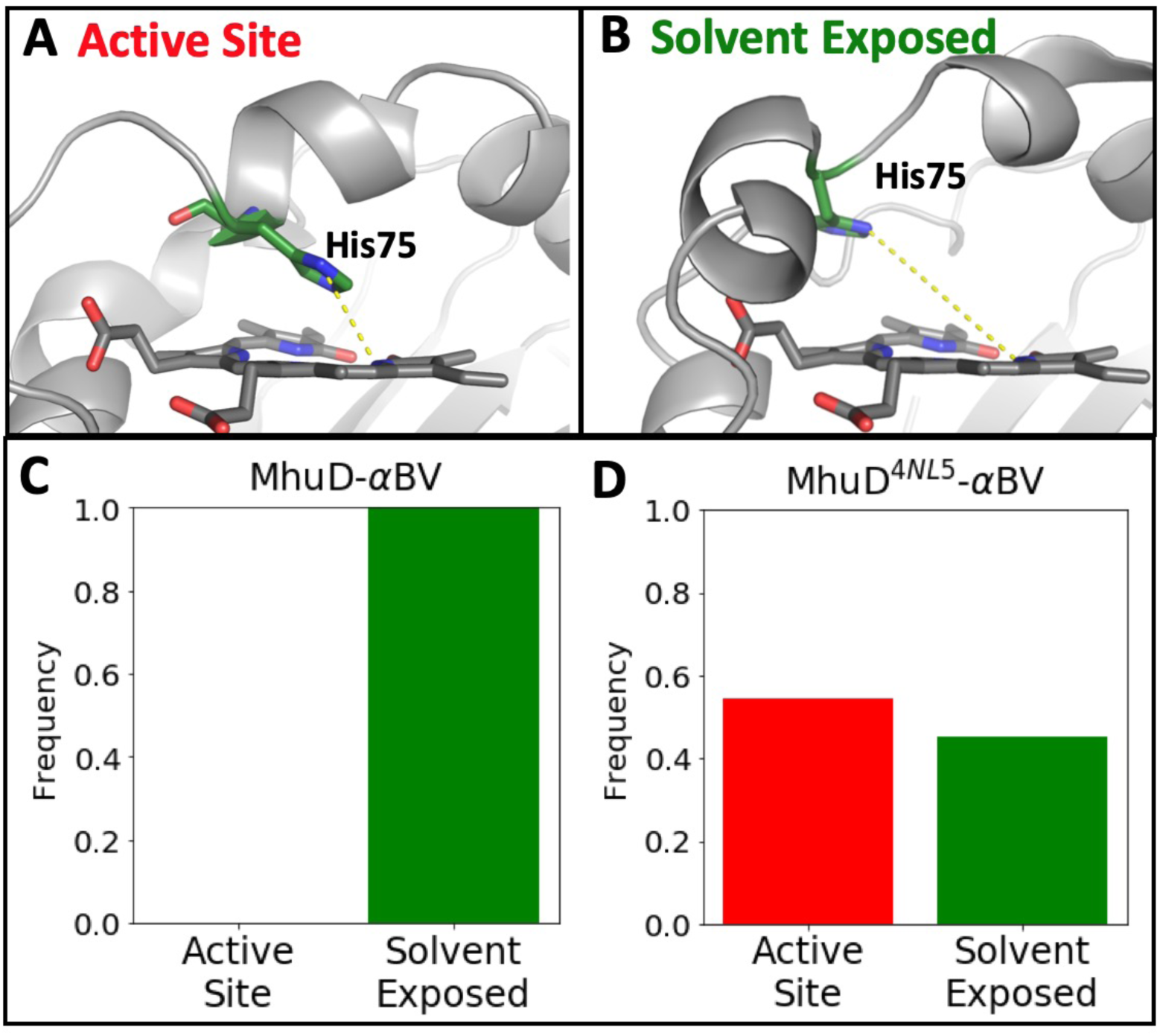
Orientation of His75 in MD simulations of MhuD. During MD simulations of MhuD with αBV, the position of His75 can be classified as either a) directed into the active site or b) rotated out of the active site. This designation was assigned based on the distance between the His75 *ε* nitrogen atom and one of the αBV nitrogens, as described in the methods section. The structure in **A.** is a snapshot from the MhuD^4NL5^-αBV simulation while **B.** shows a snapshot of the MhuD-αBV simulation. The frequency of each position is plotted for the MhuD-αBV simulations initiated from **C.** the MhuD-R26S-αBV crystal structure and **D.** the MhuD^4NL5^-αBV simulations, where αBV has docked in place of heme in the MhuD-monoheme structure. For the MhuD-αBV simulations, His75 remains oriented out of the active site while in the MD of MhuD^4NL5^-αBV, His75 flips in and out of the active site. Notably, data for MhuD^4NL5^-heme simulations of the MhuD- monoheme structure is not shown as the His75 residue is ligated to the heme-iron and thus cannot explore alternate positions.

Because the Arg79 side-chain is unresolved in the MhuD^4NL5^-heme structure but stabilized in the MhuD-αBV structure, we were also interested in exploring its dynamics in the presence of heme versus αBV. Upon inspection of the MhuD^4NL5^-heme simulations, four classifications of Arg79 positions were identified: helix 1 and helix 2 (interacting with residues on α-helix-1 or on α-helix-2), active site (interacting with heme or αBV) and solvent exposed (Figures 7A-D). In the MhuD^4NL5^-heme simulations, the varied distribution of the Arg79 positions mirrors its disorder in the crystal structure (Figure 7E). Unexpectedly, we found at times that the Arg79 forms H-bonds with the propionate groups of the heme; in fact, in the extended 600ns simulation of MhuD^4NL5^-heme, the Arg79 residue for one MhuD subunit flipped into the active site and remained coordinated to the heme ligand for over 90% of the simulation (data not shown). In simulations with αBV, Arg79 interacts with residues on helix 2 or becomes solvent exposed. Consistent with the MhuD-αBV structure, the predominant state in the MhuD-αBV simulations shows it interacting with helix 2 (Figure 7F). In our MhuD^4NL5^-αBV simulation, the favored state is solvent exposed (Figure 7G), but we suspect the interaction of Arg79 with helix 2 may be further stabilized upon full formation of α-helix-3. Given that we only observe Arg79 interacting with the ligand propionate groups in simulations where His75 is oriented into the active site, we hypothesize that Arg79 plays a critical role in promoting catalysis (when heme is present) and facilitating product egress (via formation of α-helix-3).

**Figure 7.**
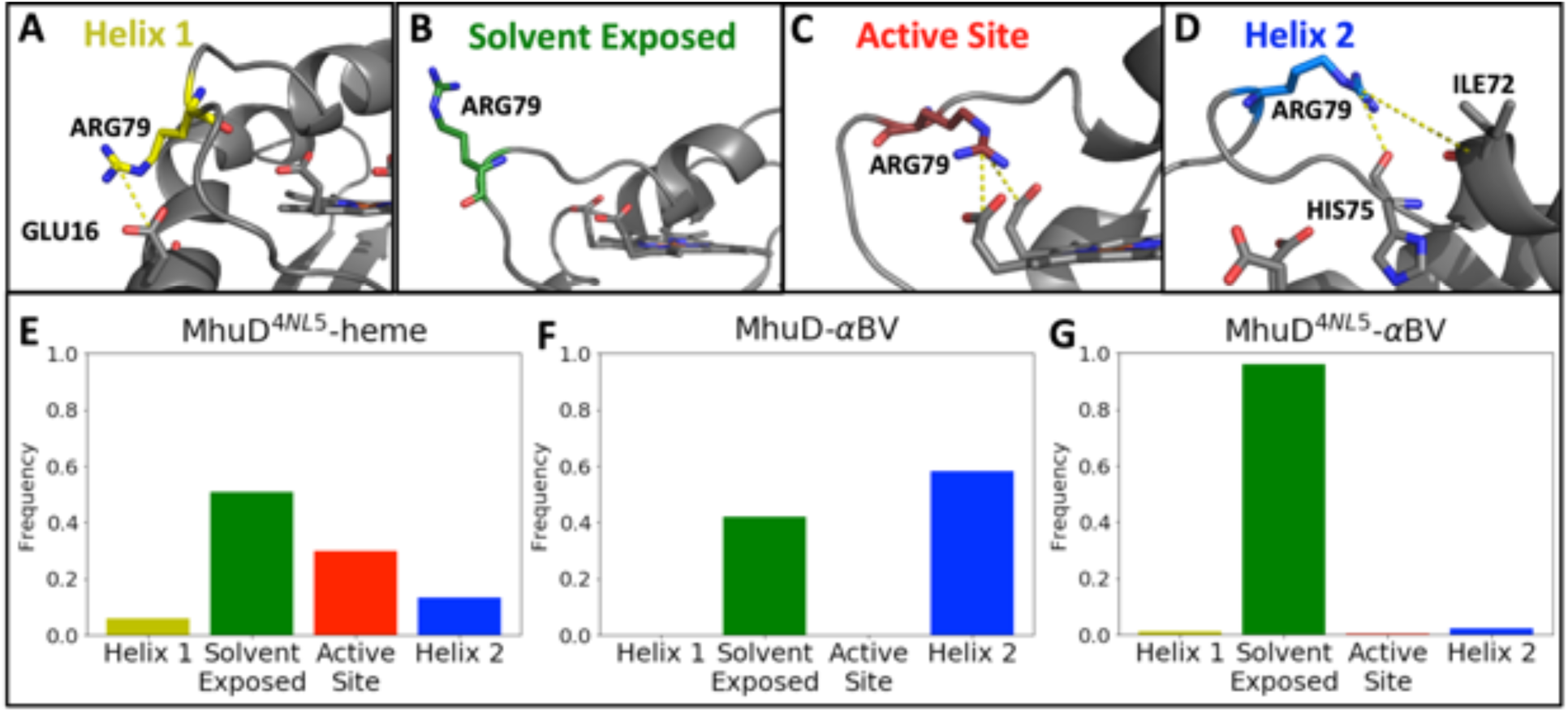
Position of Arg79 side-chain during MD simulations of MhuD. In the MhuD- monoheme structure (PDB ID 4NL5), the Arg79 side-chain is unresolved. MD simulations of the MhuD- monoheme structure (MhuD^4NL5^-heme) highlight the flexibility of this residue. We have classified its positions during simulations into four categories: **A.** *Helix 1* shows Arg79 interacting with α-helix-1 (residues 16-25), **B.** *Solvent Exposed* refers to Arg79 freely oriented in the surrounding water, **C.** *Active Site* is where Arg79 interacts with the ligand (heme or αBV), and **D.** *Helix 2* refers to the Arg79 side-chain interacting with residues in α-helix-2 (residues 60-75). All representations in **A-D** are snapshots from the simulation of MhuD^4NL5^-heme. The frequency of each Arg79 position is plotted in panels **E-G** for all three simulation types. For **E.** MhuD^4NL5^-heme, Arg79 is highly motile and at times, coordinates with the propionate groups of the heme ligand. For **F.** MhuD-αBV, where simulations are initiated from the coordinates of the MhuD-αBV structure, Arg79 predominantly interacts with helix 2 residues and may be stabilized by its participation in α-helix-3. For **G.** MhuD^4NL5^-αBV, where heme has been removed from the MhuD-monoheme structure and αBV is docked in its place, the Arg79 remains mostly solvent exposed.

Together these results suggest that in the MhuD-R26S-αBV structure we are (1) observing the proximal αBV in a location equivalent to the turnover product, (2) that the additional α-helix-3 present in the MhuD-product complex is not a crystallographic artifact and (3) the 90° rotation of His75 out of the active site is consistent with substrate turnover.

## Discussion

Within the PDB there is only one other protein structure with a strikingly similar αBV stacking conformation in its active site, a BVR from cyanobacteria (Takao et al., 2017). As with the MhuD-R26S-αBV structure, the BVR-αBV structure has offset nearly parallel tetrapyrrole planes stacked at van der Waals distance and although the proximal and distal αBV molecules in MhuD have no inter-molecular H-bonds, the BVR αBV molecules form an intermolecular H-bond between the lactam oxygen and pyrrole nitrogen, as observed between the distal and ‘bridging’ αBV molecules in the MhuD-R26S-αBV asymmetric unit (Figure 3A).

Tetrapyrrole stacking is sometimes important for enzyme activity, however it has also been observed that heme-heme stacking can be the product of crystallization. Heme stacking has previously been shown to be involved in protein electron transfer reactions, for example, the NapB, a cytochrome subunit of nitrate reductase requires two stacked heme molecules for electron transfer (Brige et al., 2002). More recently it was demonstrated that the cyanobacterial BVR required two stacked BVs to reduce BV to bilirubin (Takao et al., 2017) by an unprecedented mechanism. In contrast, MhuD can also accommodate two stacked heme molecules per active site, although this renders the enzyme inactive (Chim et al., 2010). Tetrapyrrole stacking at the crystallographic interface is not unprecedented, as observed in the structures of MhuD-diheme (Chim et al., 2010), and the ChaN-heme (Chan et al., 2006), an iron-regulated lipoprotein implicated in heme acquisition in *Campylobacter jejuni*. As MhuD is known to be active only in its monoheme form (Chim et al., 2010), and thus only produces one molecule of product per active site, we hypothesize that the proximal αBV is in the correct orientation of the MhuD tetrapyrrole product; consequently, the three additional stacked αBV molecules adjoining the two active sites of adjacent MhuD promoters are likely a product of crystallization.

The conformational changes between the MhuD substrate and product bound structures, while relatively subtle based on RMSD, are significant compared to those of canonical HOs. These differences are borne out in the fluctuation of the active site volume (Figure 8A). The structures of *C. diphtheriae* HO (HmuO) in its substrate- and product-bound form (PDB 1IW0, 4GPC) show some shifting and unwinding of the active site proximal and distal helices (Unno et al., 2013), while the change in the active site pocket volume is trivial (from ∼250 Å^3^ to ∼260 Å^3^) (Tian et al., 2018). By comparison, MhuD demonstrates much greater conformational versatility. When MhuD binds one heme molecule in its active form, the C-terminal of α-helix-1 unravels to accommodate the heme molecule and the C-terminal region of α-helix-2 extends to all the catalytically essential His75 to coordinate heme-iron, resulting in a kinked α-helix-2. The MhuD-monoheme complex can also bind another molecule of heme resulting in its diheme inactive form, whereby the kinked α-helix-2 is now extended and His75 binds to heme-iron of the solvent exposed heme molecule, nearly tripling the volume of the active site from ∼190 Å^3^ to ∼530 Å^3^ compared to the monoheme structure (Tian et al., 2018). Alternatively, when MhuD- monoheme turns over in the presence of an electron source, it forms the MhuD-product structure, which leads to the further unraveling of the C-terminal of α-helix-1 and the kinked α-helix-2 along with the formation of α-helix-3. With the transformation of substrate to product, the active site of MhuD doubles in volume, from ∼190 Å^3^ to ∼360 Å^3^ (Tian et al., 2018). The four different conformational states of MhuD highlight this protein’s inherent flexibility (Figure 8B), which may be harnessed to produce MhuD inhibitors.

**Figure 8.**
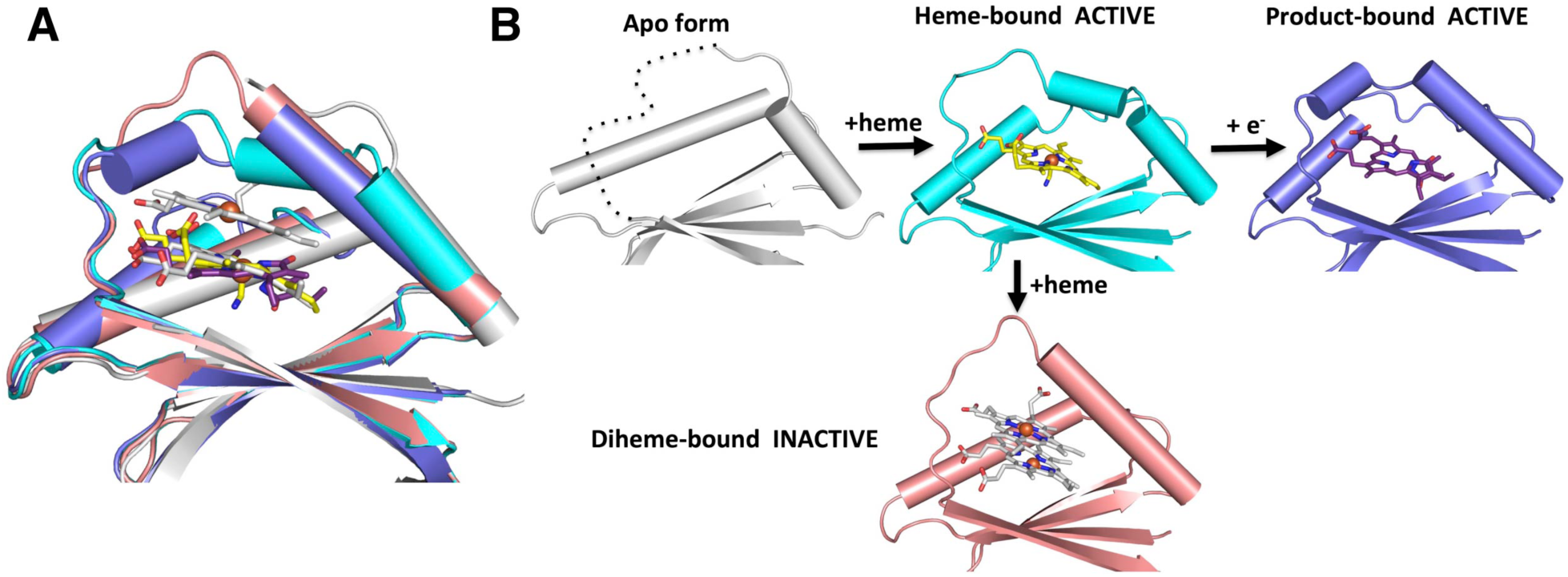
Depiction of MhuD’s conformational flexibility along its reaction pathway. **A.** Superposition of apo-MhuD (white, PDB ID: 5UQ4), MhuD-monoheme (cyan, PDB ID: 4NL5), MhuD-diheme (pink, PDB ID: 3HX9) and MhuD-R26S-αBV (blue). **B.** Apo-MhuD binds heme to form the closed MhuD-monoheme active form and in the presence of an electron donor it can degrade heme to the product bound form or alternatively, if there are high concentrations of heme, then it forms an open diheme inactive form.

Little is known about the fate of IsdG-like protein products; but removal of their tetrapyrrole products requires protein denaturation (Nambu et al., 2013; Reniere et al., 2010). Indeed, our previous and current work suggest that both MhuD substrate and product bind in the low nanomolar range and therefore the displacement of product by substrate would only occur at high heme concentrations *in vivo* (Thakuri et al., 2018). We propose three possible mechanisms of MhuD product release; (1) a dramatic conformational change would reduce product affinity and result in dissociation, (2) IsdG-type proteins are ‘suicide’ proteins that after one turnover require degradation or (3) an accessory protein is required for the removal of product from the MhuD active site. Although MhuD is an inherently flexible protein, as described above, its high affinity for tetrapyrroles decreases the likelihood that a conformational change alone would promote product release. Because it has been shown that *S. aureus* IsdG is degraded *in vivo* in its apo form, yet stabilized in the presence of heme (Reniere et al., 2011), it seems unlikely that IsdG-type enzymes are also degraded when bound to product. As the MhuD-R26S-αBV complex structure has a newly formed structural element and an associated shift in molecular surface electrostatics in comparison to MhuD-monoheme complex, these conformational changes may promote protein-protein interactions to aid in product removal.

Protein-protein interaction-induced product removal is reminiscent of human HO-1 BV removal. Human BVR interacts with hHO-1, albeit via a weak interaction, to remove the product BV and further reduce it to bilirubin (Maines and Trakshel, 1993). In contrast, the well-studied bacterial HO from *P. aeruginosa* excretes BV without further reduction (Barker et al., 2012). We hypothesize that a yet-to-be-identified Mtb protein removes product from MhuD and potentially aids in its eventual excretion, as observed in the human HO system. Surprisingly, Mtb has four close homologs of BVR even though Mtb does not have a conventional BV-producing HO enzyme (Chim et al., 2010). One of these homologs, Rv2074, has BV reduction activity although its electron donating cofactor is the flavin cofactor F420 (Ahmed et al., 2016), a deazaflavin cofactor that is a low potential hydride transfer agent (Bashiri et al., 2019), rather than flavin mononucleotide (FMN) as observed for eukaryotic BVRs (Sugishima et al., 2018). In contrast, Rv1155 does not readily reduce BV (Ahmed et al., 2015). Consequently, it was proposed that one of the BV inactive Mtb BVRs catabolizes mycobilins. Rv2607 and Rv2991 have not been tested for BVR activity and could also act in MhuD product breakdown (Ahmed et al., 2015). Mtb has both heme and siderophore-mediated iron acquisition systems (Chao et al., 2019), however *Mycobacterium leprea* only has a heme uptake and catabolism pathway. The *M. leprea* proteome only has the Mtb BVR homologs, Rv1155 and Rv2607, suggesting one of these proteins is perhaps involved in MhuD product removal.

Finally, the MhuD-product complex has an additional α-helix-3 and an accompanying change in electrostatic surface potential, compared to the MhuD-substrate complex, which may be essential for mediating protein-protein interactions to promote product removal. Notably, *S. aureus* IsdG and IsdI have a similar ^75^HisXXXArg^79^ motif encompassing the α-helix-3 (Figure 9), suggesting that both *S. aureus* IsdG and IsdI also form the new secondary structure element upon product formation; this may be a common structural feature among all IsdG-type proteins in complex with product.

**Figure 9.**
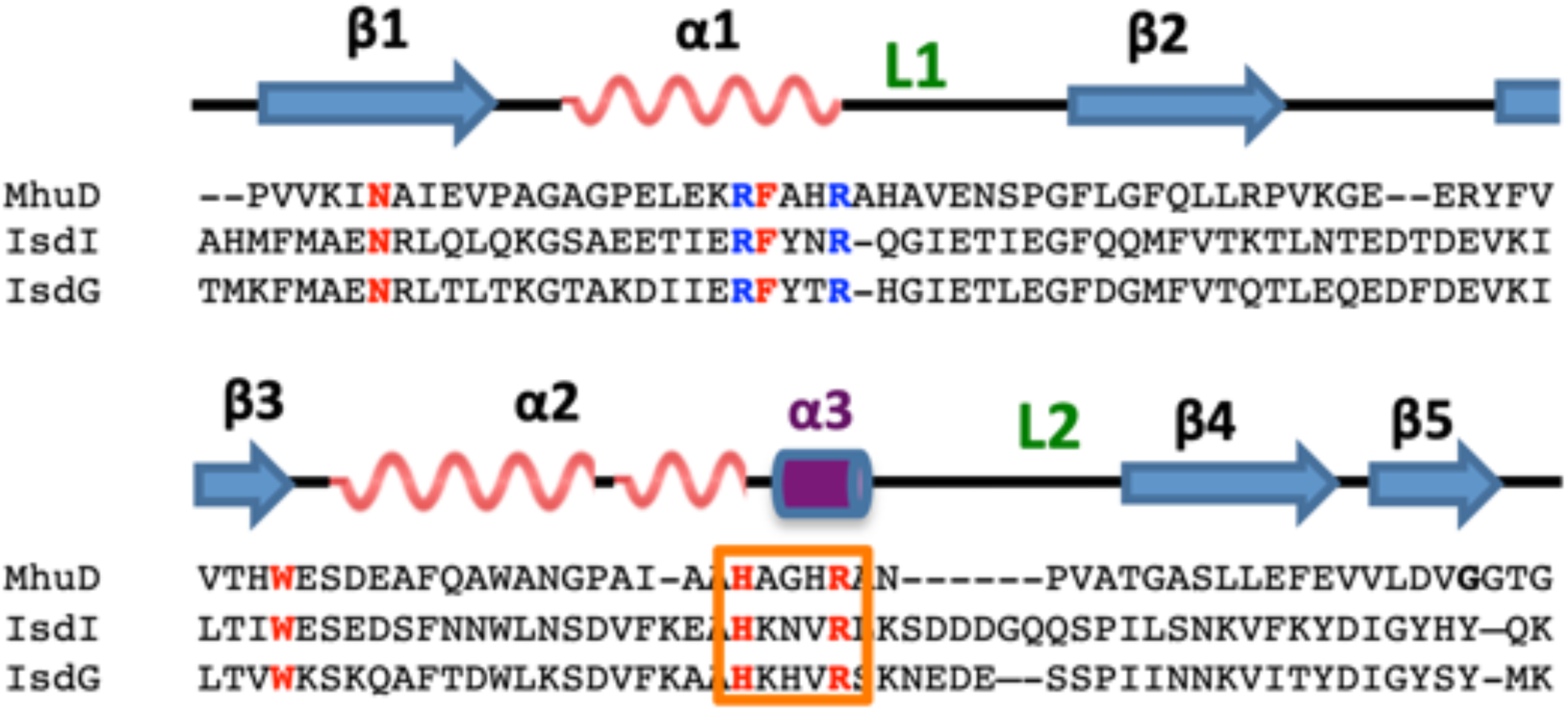
Structure- and sequence-based alignment of MhuD and *S. aureus* IsdG and IsdI. Structure-based alignment of MhuD, IsdG and IsdI. The secondary structural elements (arrows are β-strands and waves are α-helices) are based on the MhuD- monoheme structure (PDB ID: 4NL5), apart from the purple cylinder α-helix-3, that is the new helix observed in the MhuD-R26S-αBV structure. Important conserved residues are in red and blue. The MhuD ^75^HXXXR^79^ motif is highlighted by an orange box.

## Acknowledgements

We would like to thank Tom Poulos for discussions and critical reading of the manuscript, as well as Xiaorui Chen for advice on the more difficult aspects of structure refinement. We thank the Advanced Light Source at Berkeley National Laboratories (ALS) and the Stanford Synchrotron Radiation Lightsource (SSRL) for their invaluable help in data collection.

## Funding

C.W.G. thanks the National Institutes of Health (NIH) for financial support (P01- AI095208), A.C. thanks the National Science Foundation for predoctoral fellowship support (DGE-1321846), and K.H.B. thanks the NIH for support from a predoctoral training grant (T32GM108561).

## Notes

The authors declare no competing financial interests.

## Accession IDs

hHO-1 P09601

HmuO Q54AI1

MhuD P9WKH3

IsdG Q7A649

IsdI Q7A827

